# Modelling the metabolic consequences of antimicrobial exposure

**DOI:** 10.1101/2022.06.08.495414

**Authors:** Tania Alonso-Vásquez, Christopher Riccardi, Iacopo Passeri, Marco Fondi

## Abstract

Besides genetic mutations, the metabolic state of bacterial cells represents another driving factor in the emergence of antimicrobial resistance and in the actual efficacy of treatments. In this direction, studying how bacteria reprogram their metabolism when facing antimicrobial exposure is crucial to enhance our ability to limit the development and spread of antibiotic resistance. Here we have studied the metabolic consequences of antimicrobial exposure in bacteria using an integrated approach that exploits transcriptomics and computational modelling. Specifically, we asked whether common metabolic strategies emerge during the exposure to antimicrobials, regardless of the kind of antimicrobial used or, on the contrary, antimicrobial-specific pathways exist. To this purpose, we have used an heterogeneous dataset from six published studies on *Escherichia coli* exposed to different concentrations/types of compounds. We show that experimental condition, not antimicrobial exposure, is the factor that influences the most the resulting metabolic networks. However, despite condition-dependent metabolic signatures being evident, specific changes in flux distributions by antimicrobial exposed cells could be identified. In particular, purine and pyrimidine biosynthesis, and cofactor and prosthetic group biosynthesis were commonly affected by all considered antimicrobials. This suggests the presence of general metabolic strategies to face the stress posed by antimicrobial exposure and that, in turn, may represent an untapped resource for the fight against microbial infections. Finally, our analysis predicted an overall metabolic rewiring following bacteriostatic vs. bactericidal drug exposure that is in line with the current knowledge about the effects of these two classes of compounds on microbial metabolic phenotypes.

**IMPORTANCE:** A mechanistic understanding of microbial metabolic reprogramming during antimicrobial exposure is key to facilitate the discovery of new resistance mechanisms and to identify novel areas of intervention to face microbial infections. This study shows how the integration of transcriptomic data and genome-scale metabolic modelling can be used to address this critical issue, and to trace general metabolic strategies exploited by bacteria to face the stress posed by antimicrobial drugs.

---

Infectious diseases are among the top 10 causes of death worldwide. Nowadays, the most relevant factor accounting for the high incidence, morbidity and mortality of infectious diseases in developed countries is the spread of strains resistant to multiple drugs, which is leaving clinicians with very limited options to treat certain infections (1). Antibiotic resistance has become a urgent global problem, endangering the efficacy of these types of treatment for a wide range of infections. This crisis has been attributed to the overuse and misuse of them and parallelly, the lack of new drugs development (2). Contrasting antibiotic resistance is thus a priority of the public health agenda, and there is an urgent need to find novel strategies to combat antibiotic resistance. Bacterial antibiotic resistance is typically conferred by the emergence and/or acquisition of resistance genes and mostly spreads through plasmid-mediated horizontal transfer (3). However, it is becoming increasingly evident that: i) bacteria can survive under extensive antibiotic treatments without genotypic changes (4) (antibiotic tolerance (5)), and that ii) microbial metabolism plays a central role in this regard (6). This latter point has been expanded by Stokes and colleagues into three main postulates that summarise the overall relationship between bacterial metabolism and antibiotic efficacy (6). By providing a comprehensive literature survey that blends together decades of independent results, these authors conclude that i) antibiotics alter the metabolic state of bacteria, which contributes to the cellular death or stasis; ii) this resulting overall cell state influences bacterial susceptibility to antibiotics and iii) antibiotic efficacy can be enhanced by altering the metabolic state of bacteria. A further indication of the close relationship between bacterial metabolism and antibiotic resistance is represented by the observation that both drug action and resistance are global, mutually dependent perturbations of a microbial system and that they act on the cell’s metabolic network, occasionally on different components (7, 8). The interdependence of metabolism and antibiotic tolerance and the observation that the metabolic state of a bacterium is closely related to its environment including nutrients, leads to the identification of the bacterial metabolic network as a possible reservoir of novel molecular targets to fight antibiotic tolerance and, potentially, antibiotic resistance. Indeed, in many cases, the supplementation of specific nutrients has been shown to improve bactericidal activity (9, 10, 11).

Computational modelling of microbial metabolism is a powerful tool for simulating intracellular fluxes at the system level and for deriving important insights into cellular responses in specific boundary conditions. These conditions work as constraints to a genome-scale metabolic model (GSMM) in order to obtain an optimal solution of fluxes that enhances and optimises an objective function, forming a constraint-based metabolic model (12, 13) (CBMM). One of the widely used methods for the task is the Flux Balance Analysis (FBA), which calculates the flow of metabolites through a metabolic network (mass balance). FBA mathematically represents the metabolic reactions and constraints with a system of linear equations, and through the use of linear programming, it calculates the fluxes that maximise (or minimise) a given objective function (e.g. growth) previously described by those equations. The constraints help to reduce the space of feasible flux distributions (12, 14) (e.g. transcriptomic data to compute condition-specific metabolic flux changes).

Recently, researchers have started applying such techniques to frame the reciprocal interaction between cellular metabolism and antimicrobial exposure/resistance. One example is the work by Zhu et al. (15), who studied how *Pseudomonas aeruginosa* alters its metabolism, in response to polymyxin treatment. To this end, Zhu’s group developed a genome-scale metabolic model by merging two existing models, and completed the missing components using databases like KEGG (16) and MetaCyc (17), and literature. Transcriptomic data from polymyxin treatment were used as flux constraints for the developed GSMM (iPAO1), and consequently performed a single-gene deletion analysis. They showed that polymyxin treatment may reduce growth and affect a broad range of pathways, such as those related to cell envelope biogenesis. Another example of the use of GSMMs is the one of Banerjee and Raghunathan (18) who studied the case of metabolic reprogramming when *Chromobacterium violaceum*, an opportunistic human pathogen, is exposed to antibiotics. They developed the model through an initial automated draft reconstruction using the Model SEED server (19) and the *C. violaceum* genome sequence, with a further manual curation based on legacy data. The use of this GSMM (iDB858), constraint to represent three different conditions (wild type, exposure to chloramphenicol and to streptomycin, independently), described a rewired central and redox metabolism in the presence of each antibiotic, specifically, high levels of NAD recycling provided by pyruvate, suggesting a reprogramming of metabolism to compensate the stress consequences.

Finally, GSMMs can also be used to clarify if bacteria’s growth rate and metabolic state had a separate effect in antibiotic lethality. To do so, Lopatkin et al. (20) established the conditions where these two parameters were coupled and uncoupled, and used them to explore *Escherichia coli*’s response to nine different antibiotics. These data were implemented to set *E. coli*’s model (iJO1366 (21)) and perform an FBA. Their results showed that the metabolic state of bacteria correlates with antibiotic lethality, while uncoupled growth rates do not. Therefore, using metabolic simulations, they could conclude that growth-dependent effects are not sufficient to account for antibiotic-mediated lethality, but instead, the metabolic response following the initial drug target interaction showed to be crucial.

The current availability of studies addressing the short-term response to antimicrobials offers the possibility to bridge the gaps between genomics and metabolic phenotypes through the prediction of metabolic responses to various antimicrobial treatments. Here we have addressed this topic integrating genome-scale metabolic modelling with gene expression data for the model organism *E. coli*. Overall, our simulations showed trends that are consistent with the current knowledge about the intertwined relationship between antimicrobial and cellular metabolism. These included, for example, the identification of key short-term rearrangements upon antimicrobial exposure and the well-documented (opposite) metabolic phenotypes induced by bacteriostatic vs. bactericidal compounds.

## RESULTS

In this work we have focused our attention on the metabolic rewiring following antimicrobial exposure in *Escherichia coli* str. K-12 substr. MG1655. We scanned the two most important databases containing gene expression datasets (SRA and GEO) for experiments that matched the following scheme: *E. coli* grown both in presence and absence of a compound having antimicrobial activity and with transcriptomics (i.e. RNA-Seq) data obtained in each of the two conditions. All the experiments in which it was not possible to uniquely identify the exact growth conditions and/or the nature of the antimicrobial compound were disregarded for downstream analyses. This search led to the identification of six studies (22, 23, 24, 25, 26, 27) accounting for seven experiments (control vs. treatment), using as antimicrobials triclosan (TCS), colistin (CST), thioacetamide-linked 1,2,3-triazole (TAT), ampicillin (AMP), ciclopirox (CPX), erythromycin (ERY) and clindamycin (CLI), as listed in Table S1. We refer to “NAM” and “WAM” models to indicate the control (no antimicrobial) and treatment (with antimicrobial) conditions, respectively. To standardise the analysis protocol, all the reads associated with each experiment were downloaded from the public repositories and were re-analysed as described in Methods. The complete list of the experiments analysed in this work is provided as Supplementary Material (Table S1).

### Integration of RNA-Seq data with genome-scale metabolic models

Following the retrieval of RNA-Seq reads for each sample, we mapped gene expression values onto the most recent genome-scale metabolic reconstruction (iML1515) using rFAST-CORMICS (28). More in detail, for each condition, we reproduced the growth conditions used for the actual experiment by defining an *in silico* nutritional environment that resembled the one described in each work. Then, the biomass assembly reaction was selected as the objective function for each model in each condition and FBA simulations were performed in order to predict and compare the phenotype states of bacteria in WAM and NAM conditions. Figure 1A and 1B display the main features of the WAM and NAM models in terms of cumulative and antimicrobial-specific active (i.e. carrying flux) reactions. As shown, the number of active reactions in NAM and WAM models is overall similar (Figure 1A). Indeed, no significant differences were found between the number of active reactions of NAM and WAM models (T-student, p=0.23). Moreover, the number of active reactions displays a large variability in WAM models, with some of them showing a pronounced reduction in respect to the untreated (NAM) models (e.g. triclosan and erythromycin exposure), while others showing a similar (or slightly higher) percentage of active reactions in respect to their NAM counterparts (Figure 1B).

**FIG 1.**
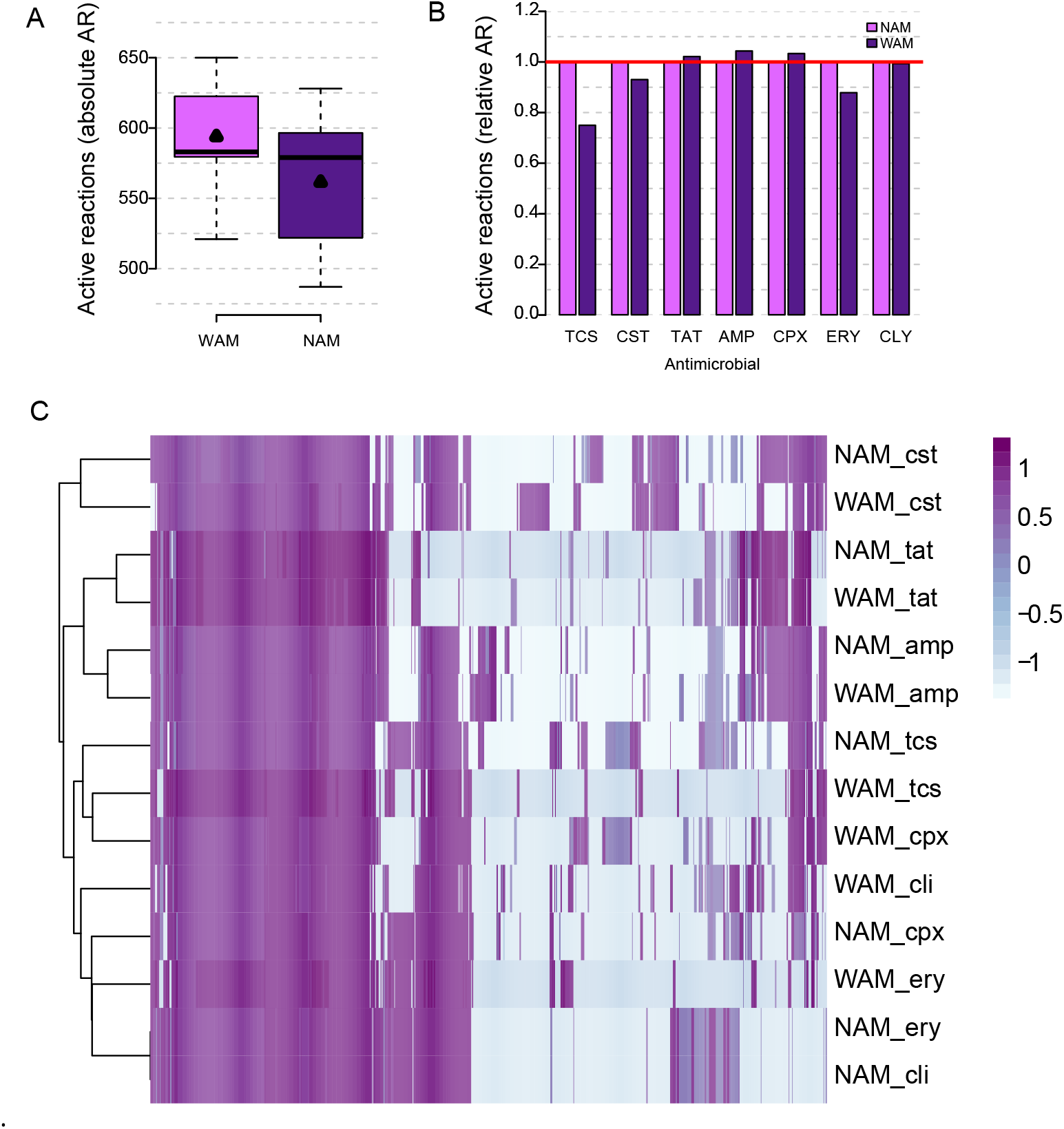
A) Number of active reactions (i.e. carrying a non-zero flux during growth simulations) in NAM and WAM models. Black triangles indicate mean values. B) Normalised active reactions (i.e. those carrying a non-zero flux during growth simulation divided by the wild type value) of NAM (dark colour) and WAM (light colour). Values lower than 1 indicate fewer active reactions in the WAM model than in the NAM model. Vice-versa, values greater than 1 indicate more active reactions in the WAM model than in the NAM model. C) Heatmap showing the clustering of NAM and WAM models (rows) in respect to the fluxes (log-normalised data) as per FBA simulations (columns).

This first set of simulations seems to suggest that the exposure to antimicrobials induces antimicrobial-specific metabolic rewiring that leads to different underlying metabolic networks, at least for what concerns the “size” (i.e. the number of active reactions) of each WAM. We then asked whether this feature (i.e. conservation of the number of active reactions) also mirrored functionally different models, i.e. whether the reactions that resulted to be active/inactive in all WAM models overlapped to some extent. In other words, we asked whether i) there exists a generalised metabolic response following antimicrobial exposure, ii) every antimicrobial elicits a specific metabolic response that results in a different metabolic network from the non-exposed one or iii) antimicrobial exposed and non-exposed models still resembles each other. According to the first scenario, one would expect all the WAM models to group together when applying a clustering algorithm that considers active reactions and corresponding experiments. The second hypothesis, instead, would imply a noisy clustering where NAM and WAM models would intermix each other. The third hypothesis, instead, would result in a pairwise clustering of WAM and NAM models. As shown in Figure 1C, our analysis depicted this latter scenario, since most of the WAM models cluster together with their NAM counterparts or in their proximity.

We then examined in deeper detail the distribution of metabolic fluxes in WAM models to identify possible common metabolic strategies in response to antimicrobial exposure. To this aim, we first identified, for each WAM model, the set of all the reactions showing a change in their fluxes when compared to its NAM counterpart and then computed the intersection of these (7) sets. Accordingly, this group of reactions may represent key metabolic pathways solicited by antimicrobial compounds exposure. On average, 532.8 (s.d. 50.3) reactions in each model displayed a flux change (increase or decrease) in respect to the respective NAM. Importantly, more than 50% of these reactions are shared by all the WAM models (Figure 2A) but not by their NAM counterparts. Thus, it appears that, regardless of the antimicrobial compound and/or its concentration and/or the overall duration of the exposure, common metabolic strategies arise.

**FIG 2.**
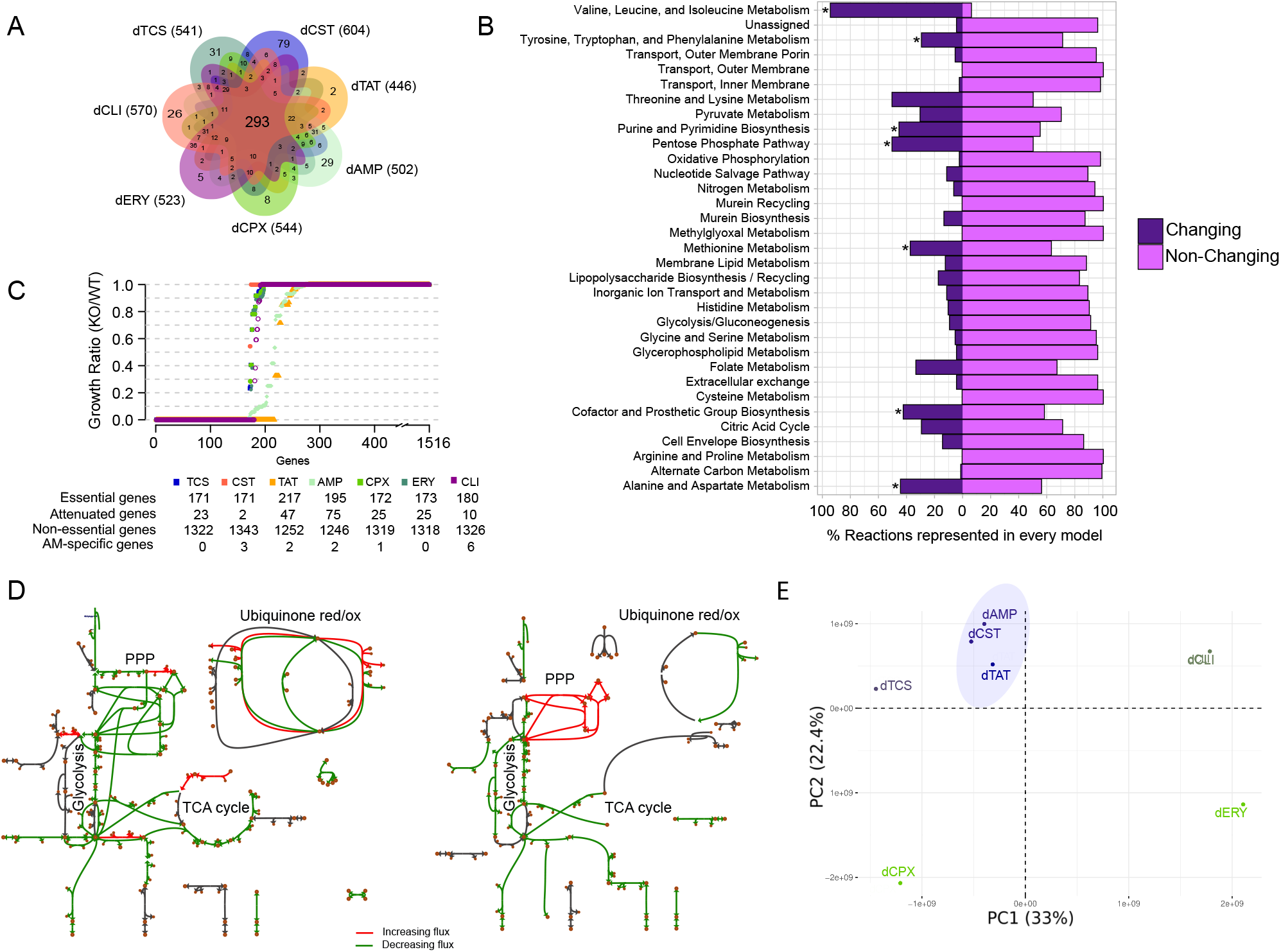
A) Venn diagram accounting for the sharing of reactions whose fluxes are altered by the exposure to the antimicrobials. B) Functional enrichment analysis of reactions changing their flux in response to antimicrobial exposure. Asterisks indicate significantly enriched functional categories (*p* − *value* < 0.05). C) Essential, attenuated (Table S4) and non-essential genes for each WAM model. Antimicrobial-specific essential genes are individuated after the gene ratio analysis. D) Schematic representation of fluxes alteration following exposure to bactericidal (left) or bacteriostatic (right) antimicrobials. Reactions represented in red/green display a consistent increase/decrease in all the models, respectively. E) Principal Component Analysis of types of antimicrobials and their reactions’ flux changes.

To functionally characterise this shared metabolic reprogramming, we performed a functional enrichment analysis based on the corresponding subsystem of each reaction in the models. Seven metabolic pathways had a significant number of changing reactions (Binomial test, *p* < 0.05) regardless of the antimicrobial treatment: methionine metabolism; valine, leucine and isoleucine metabolism; tyrosine, tryptophan and phenylalanine metabolism; purine and pyrimidine biosynthesis; pentose phosphate pathway; cofactor and prosthetic group biosynthesis; and alanine and aspar metabolism (Figure 2B).

Additionally, we focused our analysis on the identification of antimicrobial-specific essential genes (EG), that is those genes that are not required to sustain wild-type growth but become essential upon antimicrobial exposure. Hence, we systematically evaluated relevant switches in both unconstrained and constrained models, by imposing the necessary constraints to the metabolic reconstruction. Accordingly, screens for EGs were performed on each NAM and WAM pairs and WAM-specific EGs were identified. More specifically, we explored the gene essentiality (see Material and Methods) by analysing the single-gene deletion growth ratio, and found the essential, attenuated and non-essential genes related to each antimicrobial (Figure 2C). Even though we did not identify possible common essential genes in response to all antimicrobial exposure (i.e. EGs present exclusively in WAM intersection), we could list antimicrobial-specific essential genes, as described in Table S3. In accordance with flux distribution analyses (Figure 2B) our analysis highlighted 2 main pathways in which gene essentiality emerges as a consequence of antimicrobial exposure, i.e. purine and pyrimidine biosynthesis, and cofactor and prosthetic group biosynthesis. These included, for example, *purN* (Phosphoribosylglycinamide formyltransferase) for TAT exposure, *ytfE* (Iron-sulfur cluster repair) for ciclopirox resistance, and *glyA* (Alanine transaminase) for clindamycin exposure.

Furthermore, the relatively small number of WAM-specific EGs (compared to reactions changing their flux) suggests that the main short-term adaptation to antimicrobial exposure may consist in a fine-tuning of metabolic fluxes, rather than their complete rerouting to previously untapped regions of the network (i.e. the appearance of WAM-specific EGs).

### Metabolic response to bacteriostatic vs. bactericidal antimicrobials

Our dataset combines data from exposure to bacteriostatic and bactericidal antimicrobials (Table S1); the latter formed an evident cluster when performing a PCA according to the flux change of the reactions (Figure 2E). We investigated the influence on cellular metabolism of these two different types of compounds at the level of the central network, as these two classes of antimicrobials are known to elicit a profoundly different response at the metabolic level (29). In particular, we selected those reactions that consistently increased/decreased their flux in each bacteriostatic treatment and, analogously, those reactions showing increase/decrease in all bactericidal treatments (75.5% for bacteriostatics and 83.3% for bactericides). This information was then visualised using Escher (30) (Figure 2D). Overall, we noticed a general decrease in fluxes (green) both following bacteriostatic and bactericidal treatments, especially for what concerns the glycolytic region of the network. In the case of exposure to bactericidal drugs this general flux downregulation extended to the PPP (Pentose Phosphate Pathway) and of (part of) the TCA (Tricarboxylic Acids) cycle (Figure 2D left). No consistent trend could be detected for the TCA cycle of bacteriostatic exposed models (Figure 2D right), whereas the PPP of these models showed a constant flux increase throughout all the bacteriostatic treatments. Importantly, bactericidal antimicrobials activity was associated with accelerated respiration whereas exposure to bacteriostatic antimicrobials was in general associated with suppressed cellular respiration.

## DISCUSSION

In this work, we sought to identify the metabolic impact of antimicrobial exposure in microbial cells. To this aim, we have assembled (from already published studies) a highly heterogeneous dataset that comprised seven RNA-seq experiments that evaluated the transcriptomic response in *E. coli* following the exposure to this broad range of antimicrobial agents (including compounds with bactericidal or bacteriostatic activity). After a thorough re-analysis of these expression data, we integrated normalised gene counts (TPM) with genome-scale metabolic modelling to simulate how and to which extent the exposure to antimicrobials altered the fluxes inside the cell. This allowed the construction of antimicrobial exposed metabolic models (WAM) and non-exposed models (NAM) and evaluating their differences and similarities. Overall, our results indicate a marginal deviation of the WAM models from their NAM counterparts, both in terms of the overall number of flux-carrying reactions (active reactions, Figure 1A and 1B) and in terms of which reactions are turned on/off upon antimicrobial exposure (Figure 1C). Indeed, when applying a clustering algorithm to the entire set of constrained reconstructions and considering which metabolic reactions are present/absent on the basis of the expression of the corresponding genes, WAM and NAM models from the same treatment tend to be close to each other inside the resulting dendrogram (Figure 1C). Besides indicating the absence of-a generalised metabolic response following antimicrobial treatments, the similarities between WAM and NAM models highlight the influence of growth conditions (e.g. growth medium, temperature, etc.) in determining the set of differentially expressed genes and, consequently, the structure of the resulting metabolic networks. Thus, our results indicate that, in order to provide an unbiased overview on the metabolic effects of antimicrobial exposure, it is crucial to adopt the same overall growth conditions for treated and untreated cells. Ideally, the cellular response to antimicrobial treatments should be evaluated in the same experiment, with the same overall culture conditions for each condition tested.

Despite the strong similarities emerging between each couple of WAM and NAM models, when focusing on the actual metabolic rewiring and considering the activity (in terms of simulated fluxes), we could detect an entire body of metabolic reactions that were specifically perturbed (i.e. changed their fluxes) upon exposure to the antimicrobial treatment, regardless of the antimicrobial type. This set of 293 reactions (Figure 2A) represents almost one fifth of the entire *E. coli* metabolic network characterised so far, and might indicate an untapped reservoir for the elucidation of the interconnections between metabolism and antimicrobial response. Functionally speaking, these reactions were significantly associated to amino acids metabolism, purine and pyrimidine metabolism, PPP and cofactor metabolism. Interestingly, marked changes to nucleoside, nucleotide, and purine/pyrimidine base levels in response to a set of three antibiotic treatments were observed in Belenky et al. (31). Similarly, alterations of amino acids and nucleotides in response to translation and transcription inhibitors (such as some of the antimicrobials considered in this work, see Table S1) were characterised in Lobritz et al. (29), suggesting the presence of bottlenecking of metabolic flux in these pathways as a direct result of bacteriostatic/bactericidal activity. Finally, intermediates of nucleotide and amino acid biosynthesis were the most recurrently affected metabolites among different drug treatments in Zampieri et al. (32). Based on this overlap among independent studies, we showed that, despite the high heterogeneity of the assembled dataset, our method could identify pathways and biological processes involved in the global (and probably non-specific) cellular response to antimicrobial treatments. We also sought to identify, for each treatment, those genes that become essential only upon antimicrobial exposure (Table S3). Their functional analysis showed that they overall participate in the same pathways that embedded the higher number of changing reactions (i.e. purine and pyrimidine biosynthesis).The relatively small number of antimicrobial-specific essential genes, however, suggests that the primary response to antimicrobial treatments probably involves a general metabolic reprogramming (many reactions changing their activity in respect to the non-exposed metabolic network), rather than the activation of previously inactive metabolic routes (the appearance of essential genes only in the exposed metabolic network). Nevertheless, the body of essential and attenuated genes identified here may provide new targets to improve the efficacy of antimicrobial treatments leveraging on pathogens metabolism (9, 10, 11).

Metabolically speaking, one of the most striking features of antimicrobial treatments is the different impact caused by bacteriostatic vs. bactericidal drugs. Many studies (31, 29, 33, 34) have shown that, while growth inhibition by bacteriostatic compounds is typically associated with diminished cellular respiration, bactericidal antibiotics action usually induces an accelerated respiration. We thus focused our analysis on the central metabolism of *E. coli* and predicted the metabolic response through FBA following the exposure to bacteriostatic and bactericidal treatments. As shown in Figure 2D, in both cases a general downregulation of metabolic activity (green fluxes) was observed, in agreement with Chen et al. (35). However, significant differences emerged in the comparison between bacteriostatic vs. bactericidal induced metabolic networks. In particular, we observed a consistent increase in the respiration rates of all bactericidal treatments whereas the bacteriostatic treatments all induced lower respiration rates (Figure 2D). This finding further confirms that bactericidal and bacteriostatic compounds induce different and complex sets of metabolic changes in bacteria that ultimately translate into different fluxes distributions at the level of key, central pathways.

Interestingly, the metabolic response to the bacteriostatic compounds used in these experiments, seems to induce an increase in the fluxes of PPP (Figure 2D). PPP is a fundamental pathway in bacterial metabolism for the maintenance of carbon homoeostasis, the supply of precursors for nucleotide and amino acid biosynthesis and for the production of reducing molecules. This latter aspect has been found to be important in the exposure to bactericidal compounds that, as our study suggests, leads to the increase of the overall cellular respiration rate that, in turn, may favour the occurrence of ROS (Reactive Oxygen Species) inside the cell. In our case, the overall increase in PPP fluxes following bacteriostatic exposure may rather reflect the rerouting of the glycolytic flux into the pentose phosphate pathway. Indeed, the increase in PPP fluxes might reflect the requirement of nucleotide biosynthetic intermediates. Indeed, 1) nucleotide biosynthesis resulted to be one of the pathways that significantly included reactions increasing their fluxes (Figure 2B) and 2) previous studies, reported that gene expression changes in nucleotide biosynthesis were one of the most pronounced transcriptional responses upon different stress conditions (32).

Exposure to antimicrobials has a deep impact on bacterial physiology. Metabolism has been shown to undergo drastic rearrangements once cells are treated with antimicrobials (36). The study of such changes in cellular physiology is particularly important from both a basic research perspective as well from an applicative viewpoint. Indeed, metabolism may represent a still poorly explored reservoir of novel targets for fighting infections. Additionally, identifying those reactions/pathways that are mostly affected by such treatments may reveal those regions of the metabolic network whose elicitation may boost up the efficacy of antimicrobials. The combination of a very heterogeneous dataset of transcriptomic reads from seven different contrasts (treated vs. untreated) with genome-scale metabolic modelling has confirmed the involvement of specific pathways in the short-term response to such stressors. Our results agree with current knowledge about the intertwined relationship between antimicrobial and cellular metabolism. These included, for example, the identification of key short-term rearrangements upon antimicrobial exposure and the well-known metabolic differences induced in the cells by bacteriostatic vs. bactericidal compounds. Nevertheless, we observed that the resulting metabolic networks of WAM models systematically resembled those of their NAM counterparts. This fact suggests that the experimental conditions of such studies may have a deep impact in determining the transcriptomic background of the cell and, consequently, their presumed active metabolic network. Such dataset heterogeneity may have a dual impact on our conclusions. On the one hand, those common features listed above are probably robust as they emerged over the noisy background of such a sparse dataset; on the other hand, it may represent a critical point of this study as it may hide further connections between the tested antimicrobial compounds and their associated metabolic response. We thus highlight the importance of standardised and controlled settings when testing to rule out the possible influence of growth/experimental conditions in the final interpretation of the results.

## MATERIALS AND METHODS

### Dataset assembly and validation

We used a set of available transcriptomic data for the bacterium *Escherichia coli* retrieved from the SRA and/or GEO databases. In our experience, metadata associated with public sequencing projects (including the name of the microbial strain associated with the dataset) are often poorly curated. For this reason, we conducted a pristine quality check on every RNA-Seq sample to determine the reliability of the transcriptomic signal for the herby presented work. All samples were assessed for base call quality and adapter content using fastp (37) version 0.22.0, allowing down to a mean quality threshold of 20 (i.e., probability of incorrect base call of 1 in 100) and minimum read length of 40 nucleotides. Subsequently, we utilised Squid (38) with its default options to assess the proportion of reads that mapped to the coding sequences. We set a threshold of 80% of reads mapping to determine the samples that were to be kept. This allowed the definition of the actual dataset consisting of contrasts (treated vs. untreated). This information is detailed in Table S1.

### NGS reads processing

We used Salmon (39) to quantify the transcripts for each condition. The reference gene annotation used for this aim was the one of ASM584v2 genome. Out of this process, we obtained the TPM (transcripts per million) for each gene in each contrast.

### Genome-scale modelling

To perform genome-scale modelling, we used the latest metabolic reconstruction available for *E. coli*, i.e. iML1515 (40). rFASTCORMICS was used to constrain the models using available gene expression data. This MATLAB code-based algorithm written by Pacheco et al. (28), performs the reconstruction of context-specific metabolic models based on RNA-Seq data without a priori defined expression thresholds. This tool returned metabolic models for each treatment and their controls. These models were further constrained as follows. In each of the simulations we set the formation of biomass as the objective function and set the boundary constraints of the models to recapitulate the nutrients availability of each experiment (Table S2). All the metabolic simulations were performed using COBRA toolbox (41) in MATLAB 2020a. Context-specific models (i.e. WAM and NAM models) are available as Supplemental Material SM1.

### Identification of reactions changing in WAM vs NAM

We identified all the reactions carrying a non-zero flux for all models, i.e. all the reactions that would be active in at least one of them. We then calculated the flux ratio by dividing each WAM reaction flux by its NAM’s analogous, obtaining the change ratio for each reaction, and the set of changing reactions after the antimicrobial exposure (change ratio different from one). From these data we could calculate the set of reactions that are changing regardless of the antimicrobial used, by intersecting each individual set.

### Functional enrichment analysis

To analyse the functional enrichment, we found the number of reactions changing for each metabolic pathway (subsystem). In this regard, we assessed if the amount of changing reactions per subsystem was significant with respect to the non-changing ones using a binomial test with a significance threshold of 0.05. We performed the functional enrichment analysis only with the subsystems formed by nine or more reactions in total.

### Gene essentiality analysis

For the gene essentiality analysis, we performed an *in silico* single-gene deletion using COBRA toolbox (41), and the resulting NAM and WAM models for each condition (constrained with transcriptomic data). Genes with a growth ratio (KO/WT) lower than 0.1 were considered as essential for cell growth; otherwise, they were considered attenuated genes (0.1 < growth ratio < 0.99) or nonessential genes (growth ratio > 0.99). We looked for common essential and attenuated genes regardless of the antimicrobial used by intersecting the NAM and WAM common genes. To identify the antimicrobial-specific essential genes, we intersect the essential genes of WAM and its counterpart NAM models for each treatment.

### Identification of bactericidal and bacteriostatic key reactions

We performed a Principal Component Analysis (PCA) to assess if the exposure to bacteriostatic and bactericidal antimicrobials could explain the fluxes’ behaviour. Using these two types of antimicrobials, we formed two different sets, and in order to represent them, we selected the reactions whose fluxes were included in the space delimited by one standard deviation. These reactions were then discretized and mapped using Escher (30), to visualise them into their metabolic pathways.

## SUPPLEMENTAL MATERIAL

Supplemental material is available online only.

**TABLE S1**, XLSX file, 0.02 MB

**TABLE S2**, XLSX file, 0.02 MB

**TABLE S3**, XLSX file, 0.01 MB

**TABLE S4**, XLSX file, 0.01 MB

**FILE SM1**, Constrained Models, Dropbox folder

## ACKNOWLEDGMENTS

This work was financially supported by a PRIN (Programmi di Ricerca Scientifica di Rilevante Interesse Nazionale) grant to Marco Fondi (Escaping the ESKAPEs: integrated pipelines for new antibacterial drug, 20208LLXEJ_002).

Conceptualization, Marco Fondi; Data curation, Christopher Riccardi and Iacopo Passeri; Formal analysis, Tania Alonso-Vásquez; Investigation, Tania Alonso-Vásquez, Iacopo Passeri and Marco Fondi; Methodology,Tania Alonso-Vásquez and Marco Fondi; Project administration, Marco Fondi; Visualization, Tania Alonso-Vásquez and Marco Fondi; Writing–original draft, Tania Alonso-Vásquez and Marco Fondi; Writing–review, Tania Alonso-Vásquez, Christopher Riccardi, Iacopo Passeri and Marco Fondi.

